# Exploiting modal demultiplexing properties of tapered optical fibers for tailored optogenetic stimulation

**DOI:** 10.1101/199273

**Authors:** Marco Pisanello, Filippo Pisano, Leonardo Sileo, Emanuela Maglie, Elisa Bellistri, Barbara Spagnolo, Gil Mandelbaum, Bernardo L. Sabatini, Massimo De Vittorio, Ferruccio Pisanello

**Author notes:** These authors contributed equally to this work (Marco Pisanello, Filippo Pisano).

## Abstract

Optogenetic control of neural activity in deep brain regions requires precise and flexible light delivery with non-invasive devices. To this end, Tapered Optical Fibers (TFs) represent a minimally-invasive tool that can deliver light over either large brain volumes or spatially confined subregions. This work links the emission properties of TFs with the modal content injected into the fiber, finding that the maximum transversal propagation constant (k_t_) and the total number of guided modes sustained by the waveguide are key parameters for engineering the mode demultiplexing properties of TFs. Intrinsic features of the optical fiber (numerical aperture and core/cladding diameter) define the optically active segment of the taper (up to ∼3mm), along which a linear relation between the propagating set of k_t_ values and the emission position exists. These site-selective light-delivery properties are preserved at multiple wavelengths, further extending the range of applications expected for tapered fibers for optical control of neural activity.

## 1. Introduction

Over the past decade, optogenetic stimulation has had a remarkable impact on neuroscience research as it permits millisecond precision manipulation of genetically-targeted neural populations ^1,2,3^. However, harnessing the full potential of the optogenetic approach requires light to be delivered to the opsin-transfected neuron under tight spatiotemporal control. This task is even more arduous when targeting brain regions beyond 1 mm depth, as common light-delivery methods (cleaved fiber optics and two-photon excitation) fail in penetrating such depths or are highly invasive. Driven by the experimental need for novel approaches, research in innovative technologies for deep brain light delivery has produced several promising solutions such as multi-dimensional wave-guides or high-density μLED probes^4–20^. In a recent work^21^ we demonstrated a simple and cost-effective device based on a thin Tapered Optical Fiber (TF) that can perform both homogeneous light delivery over large volumes and dynamically–controlled spatially-restricted illumination. This was done by exploiting TFs light-emission properties that are determined by the subset of guided modes that propagates in the taper ^22^. As modes of decreasing order and lower transversal propagation constants k_t_ are progressively out-coupled while travelling in the tapered region, light output can be modulated over a long segment of the taper surface. Experiments from two independent research groups have shown in the last year the great advantages of TFs for *in vivo* control of neural activity in different animal models. For example, wide-volume illumination was obtained in the motor cortex and in the striatum of both free-moving and head restrained mice^21^, as well as in the Frontal Eye Field of non-human primates^23^. TFs also allowed for site selective light delivery in the striatum to control specific locomotion behavior, on the base of different modal subsets injected into the fiber to illuminate ventral or dorsal striatum using only one and minimally invasive implant^21^. However, to the best of our knowledge, no studies directly link the modal subsets guided into the fiber with the light delivery geometry from a tapered optical fiber for optogenetics applications, and no direct assessment of TFs site-selective light-delivery performances at longer wavelengths has been reported, a crucial aspect for using this technology with a broad set of opsins^24–26^.

Building on this background, here we analyze the relationships between guided modes and emission properties of TFs with different numerical apertures, finding that the maximum transversal propagation constant k_t_ and the total number of guided modes sustained by the TFs are key parameters to tailor taper emission properties with the brain structure of interest. By combining far field analysis and fluorescence measurements of emitted light, we retrieve the characteristic curve linking the injected modal subset with its out-coupling position along the taper, showing that the position of the light emitting region in high-NA TFs is linearly dependent on the light input angle for both blue and yellow light.

## 2. Results

### 2.1 Effects of NA and maximum transverse propagation constant on the emission properties of tapered optical fibers

When injecting light in the whole fiber numerical aperture (NA) with a Gaussian beam, light-delivery devices based on TFs, schematically displayed in Figure 1a, can illuminate large brain volumes by emitting radiation from a long segment of the taper surface^21,23^. This occurs because of a gradual loss of light along the taper generated by the progressive narrowing of the waveguide diameter *a(z)*. Given a fiber with initial diameter *a*_*0*_, the transversal propagation constants k_t_ of the guided modes increases from the initial value k_t_(a_0_) following the relation^27^

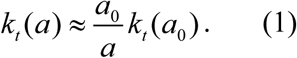

**Figure 1:**
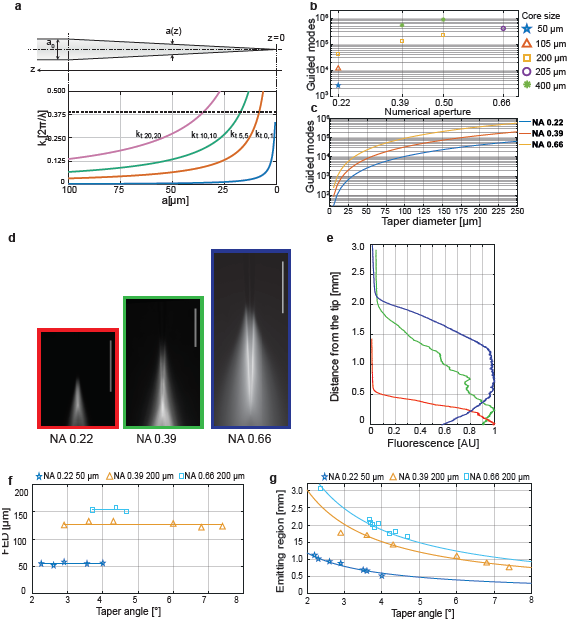
Tailoring TF devices emission lengths. (**a**) *(top)* Schematic representation of a tapered optical fiber. *(bottom)* Evolution of the transversal propagation constant of four different guided modes as a function of the taper diameter. **(b)** Number of modes sustained by fiber optics with increasing core size and NA. (**c**) Number of guided modes at increasing diameters of the tapered section for fibers with 50μm core/0.22 NA (blue curve), 200μm core/0.39 NA (red curve), and 200μm core/0.66 NA (yellow curve). (**d**) Fluorescence image of light emission for a fiber with 50μm core/0.22 NA *(left)*, 200μm core/0.39 BA *(center)*, and 200 μm core/0.66 NA *(right)*, ∼3.5° taper angle; scale bars are 1 mm. **(e)** Normalized intensity profile measured along the taper surface for the three fibers in panel (d). **(f,g)** First Emission Diameters and Emission Lengths, respectively, for different fiber types with increasing taper angle. For panels d-g data pertaining 0.22 NA and 0.39 NA fibers were reused from Pisanello et al**^21^**.

Since the taper can support light propagation only for modes with^27^

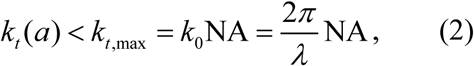

where λ is the propagating wavelength, each mode that does not fulfill this constraint is out-coupled. If the radiation injected into the tapered region of the waveguide is composed by guided modes with *k*_*t*_ in the interval [0, k_t,max_], the condition in equation 2 will be gradually broken along the taper. This is shown in figure 1a where transverse propagation constants (colored lines) overtake the k_t,max_ threshold (dashed line). As a consequence, higher order modes with higher transversal propagation constants tend to be out-coupled at larger taper diameters. Low order modes with low *k*_*t*_ are instead outcoupled close to the taper tip. It follows that an optical fiber supporting a larger *k*_*t,max*_ and sustaining a higher number of modes is expected to produce tapered sections with a longer region able to optically interface with the environment. Following Snyder et al.^27^, the total number of guided modes (*M(z)*) for a section of diameter *a(z)* can be approximated as

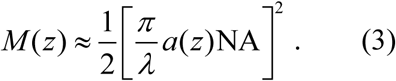

As a direct consequence, fibers with higher core size and higher NA can sustain a larger number of guided modes (Figure 1b) and the number of guided modes diminishes when approaching the fiber tip (Figure 1c). Building on this property, we selected three optical fibers that support an increasing number of modes distributed over widening intervals of transverse constants [0, k_t,max_] with similar taper angles (ψ∼3.6°). Namely, we used fibers with NA=0.22 and *a*_*0*_=50 μm, NA=0.39 and *a*_*0*_=200 μm, and NA=0.66 with *a*_*0*_=200 μm. The extent of the taper segment emitting light (hereafter referred to as “emitting length”, EL) was measured as the distance from the tip at which the detected fluorescence decreases to half of the intensity at the fiber tip (see emission profiles displayed in figure 1d,e and detailed definition in Methods). For the same taper angle (ψ∼3.6°), TFs with higher NA are optically active over a longer taper region (Figure 1e), as relation 2 starts to be broken farther from the taper tip. In detail, 0.22 NA fibers show EL∼484μm, for 0.39 NA TFs EL∼1220μm, and EL∼1797μm for 0.66 NA. Data pertaining 0.22 NA and 0.39 NA fibers were reused from Pisanello et al^21^.

Although the number of guided modes decreases as soon as the waveguide narrows (Figure 1c) and relation (2) is broken just after the taper entrance for modes with high k_t_ (figure 1a), light emission starts to be appreciable in the fluorescence measurements only at a First Emission Diameter (FED) that depends on fiber NA and waveguide size (Figure 1f). This can be ascribed to the fact that high order modes have higher propagation losses with respect to low order modes and they are excited with lower power efficiency by a Gaussian beam focused on the fiber core^27,28^. As shown in figure 1f, fibers with large core size and large NA (e.g. supporting a larger number of modes and a higher k_t,max_) lead to higher values of the FED, which do not depend on the taper angle ψ. For the fibers investigated here, this leads to the observation that the higher the NA, the higher the ratio between the FED and the fiber diameter at the taper entrance: 55/125=0.44 for NA=0.22, 126/225=0.56 for NA=0.39 and 154/230=0.67 for NA=0.66. Data pertaining 0.22 NA and 0.39 NA fibers were reused from Pisanello et al^21^. Therefore, 0.66NA fibers are expected to support for the longest ELs.

Given the independence of FED on ψ, EL is expected to depend on the taper angle for a linear taper profile following the relation

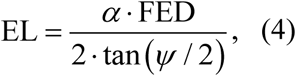

as shown by the good agreement with experimental data displayed Figure 1g. The scale factor α was defined as the ratio between the EL and the FED distance from the taper tip for ψ=3.7°, and was similar across NAs (α_NA=0.66_=0.8652, α_NA=0.39_=0.8956 and α_NA=0.39_=0.8848). The fiber allowing for the longest emission for the same ψ is the one that distributes the widest range of k_t_ on the widest number of guided modes. In light of these results, the FED can be identified as a chief design parameter for TFs, taking into account the effects on taper emission of several constitutive parameters of the fiber: the number of guided modes that propagates in the taper and of their k_t_ values in relation with k_t,max_.

These constraints can be considered to tailor the emission length of TFs to the size of targeted structures in the mouse brain, as schematically shown on the right of Figure 1g. For example, minimally invasive probes can be engineered from NA=0.22 and a=50μm fibers that can be tailored to uniformly illuminate structures ranging from ∼ 400 μm to ∼750 μm. These probes, for example, offer a promising tool to interface with deep brain regions, such as the amygdala. At the same time, a waveguide with 0.39 NA and core diameter *a*=200 μm can perform dynamically controlled light delivery over large brain volumes, ∼ 1.4 mm in the dorso-ventral direction, as recently demonstrated^21^. Finally, 0.66 NA TFs present a long and broadly-tunable emitting region that can extend from ∼ 1.5 mm up to ∼3 mm. This property might find applications in optogenetic stimulation of extended brain areas such as the whole dorsoventral extent of the mouse striatum or the depth of primate cortex.

### 2.2 Linear control of TFs emission regions

As previously demonstrated, the light emitting portion of TFs can be dynamically controlled by remotely adjusting the light input angle ^21^. In fact, coupling light into TFs with an angle-selective launching system selects well-defined subsets of bounded modes that propagate into the optical fiber, which are then out-coupled at different taper sections^21^. In the following the relationship between the input angle and the out-coupling position along the taper is estimated.

To this purpose the light redirection system displayed in Figure 2a was implemented: a lens L1 focuses a Gaussian laser beam onto the rotation axis of a galvanometer mirror (GM). The beam is deflected by GM of an angle θ_GM_ and is collimated by the lens L2 and focused onto the fiber by the lens L3 at an angle θ_in_. The taper was submerged in a PBS:fluorescein bath to analyse the light emission geometry as a function of θ_in_. 0.39 NA and 0.66 NA TFs with 200 μm core diameter were tested, both with ψ∼3.5°. As already mentioned in the previous paragraph, 0.66 NA fibers sustain a high number of guided modes (∼4.0×10^5^) distributed over a large ensemble of transverse propagation constants (k_t,max_=8.7×10^−3^ nm^−1^ for λ=473 nm). Typical images of light emission for four different θ_in_ are displayed in figure 2b. As shown by the emission profiles in the proximity of the linear taper surface (figure 2b), it is possible to dynamically redirect light output over four different emission regions with equally spaced emission peaks. By extracting the centroid of the emission region (*c*) and the relative FWHM (*Δc*) from the emission profile at each input angle (definitions of *c* and *Δc* in Methods), we found that *c* depends linearly on the input angle θ for θ>10° (with a slope of 50.89 μm/°, fit RMSE of 11μm) and can be dynamically moved along ∼1.8 mm (figure 2c). Moreover, the FWHM was found to be approximately constant at *Δc* = 550 +/−5 μm for θ>10°. This behaviour was confirmed by ray tracing simulations that, as displayed in figure 2d, highlight the linear relation between the position of the emission region and θ. The slopes of the linear fits on the centroid position for experimental and simulated data are in good agreement (50.89 μm/° vs 58.01 μm/°). However, simulations data predict a positive shift of ∼200μm with respect to experimental data. This effect is likely due to an oversimplified modeling of the refractive index of the tapered region, which was assumed to be homogeneous. For 0.39 NA fibers with the same taper angle, supporting a lower number of modes (∼1.3x10^5^ at 473nm) and a narrower range of k_t_ (k_t,max_=5.2×10^−3^ nm^−1^ for λ=473 nm), *c* can be moved along ∼1.4 mm, although the position of the centroid slightly deviates from a linear dependence (slope fit of 50.15 μm/°, fit RMSE of 76 μm), as shown in Figure 2e.

**Figure 2:**
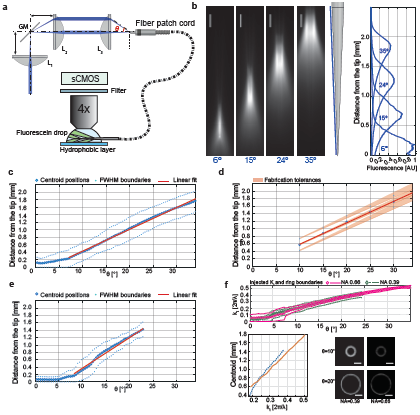
Spatially selective light delivery using high-NA TF. (**a**) Diagram of the optical setup (L1, L2, L3 lenses; GM galvanometric mirror). Laser light enters the fiber patch cord at an angle and is outcoupled from the taper surface immersed in fluorescein. (**b**) *Left*, emission profiles for a NA=0.66, 200 μm core size fiber with increasing input angle. Scale bars are 250 μm. *Center*, diagram of the tapered section. The blue line indicates the profile along which the intensity was measured. *Right*, intensity profiles measured at different input angles with respect to the distance from the tip. Four selected regions are selected by varying the input angle. (**c**) Position of the intensity centroid of the emission region (blue circles) and positions of the FWHM values of the emission region (pale blue dots) with respect to the input angle for a NA=0.66 TF. The red line indicates a linear fit performed on the centroid position. (**d**) As in panel (**c**) for ray tracing simulations; shaded area indicates fabrication tolerances. (**e**) As in panel (**f**) for a NA=0.39 TFs. (**f**) Injected k_t_ versus NA for 0.39 and 0.66 TFs. *Top*, centroid and boundaries of the injected kt- subset versus input angle. *Bottom left,* centroid position versus the injected kt for 0.66 NA, orange line, and 0.39, dashed blue line, TFs. *Bottom right,* typical of images of the patch cord far field at θ=10°, 20°. Scale bars are 0.15·2π/λ.

To confirm that the observed difference between geometrically identical 0.39 NA and 0.66NA TFs is related to the different extent of k_t_ values sustained by the two fibers, the method proposed by Pisanello et al^22^ was used to measure the subsets of *k*_*t*_ injected into 0.66 NA and 0.39 NA TFs for each input angle. Briefly, we injected light on a patch cord fiber and then imaged far field of the fiber emission on a CCD camera (supplementary figure 1). As shown in figure 2f, this produced light rings whose diameter is proportional to the input angle and can be linked to the transverse propagation constant k_t_. In order to characterize the injection of modal subset with a k_t_(θ)±Δk_t_(θ) curve, we extracted the output ring diameter and its FWHM for each input angle (see Methods). The results of the measurement are reported in figure 2g. As expected, k_t_(θ) values for 0.39 and 0.66 NA fibers are almost overlapping for θ>10°. For the same injected k_t_ value, emission from the 0.66 NA TF is closer to the fiber tip (figure 2f) due to the higher refractive index, and light emission can be moved on a wider taper extent by virtue of the higher k_t,max_ supported by the 0.66 NA TF.

Another important aspect to evaluate the influence of injected k_t_ values on TFs light emission properties is the different attenuation of light injected at different θ_in_ as high order modes have higher attenuation constant than do low order modes^27,28^. This is visible in the measurements displayed in figure 3a (red curve): before entering the taper, light power slightly decreases as a function of θ_in_ and the same happens at the taper output (back curve). Interestingly, in the range of input angle for which *c* depends linearly on θ_in_ (i.e. θ_in_ >10°), the coupling efficiency between the taper input and the taper output remains approximately constant, as shown in figure 3b. A drop in coupling efficiency was observed for low injection angles. This effect was explained by SEM inspection, which revealed lower quality of surface at the very tip of the taper (figure 3c). On the basis of these considerations, the output power density distribution around the taper for different input angles was assessed from the measured emission profiles (see Methods for details on the performed calculations). Sample images for three different θ_in_ in the linear range are shown in figure 3d. The average power density can be maintained constant over the different emission regions by slightly increasing the total output power as a function of θ_in_, compensating for the increase of the optically active surface of emission regions at higher diameters. For the particular case shown in figure 3d, an increase of ∼3 fold, roughly similar to the increase in optically active surface, holds the average power density constant (12 mW/mm^2^) for θ_in_=15°, 24°, 35°.

**Figure 3:**
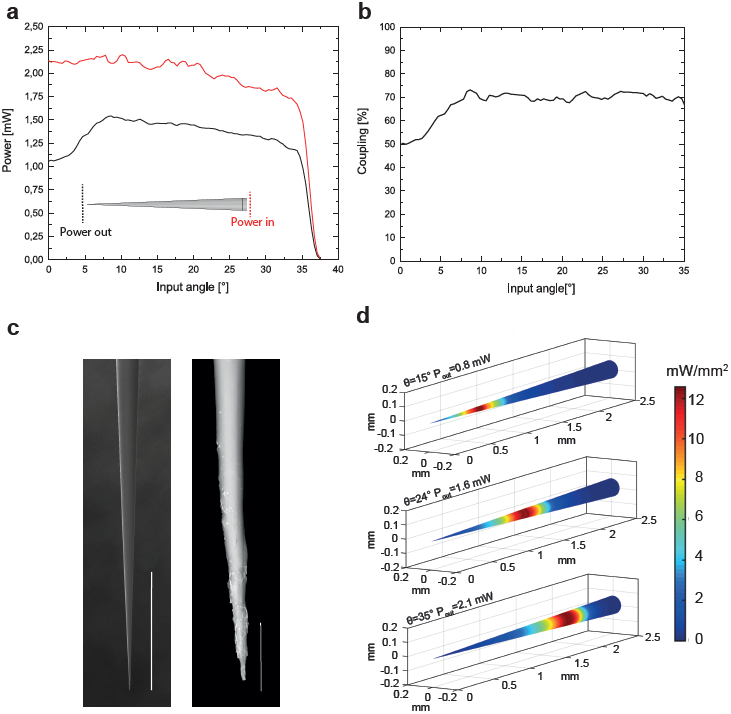
Characterization of light coupling efficiency and output power density for NA=0.66 TFs. (**a**) Light coupling across the angular acceptance. The input power (red line) was measured at the entrance of the tapered region. The output power (black line) was measures in close proximity to the taper tip, as shown in the inset. A drop in power output was observed for low input angles. (**b**) Power coupling efficiency, calculated as the ration between output and input power, versus light input angle. The coupling efficiency is approximately constant for θ_in_>10°. (**c**) Scanning electron microscope images of a NA 0.66 TF. While the left panel shows a homogeneous portion of the taper region, scale bar 400 μm, the close-up in the right panel shows degradation of the taper surface in proximity of the tip, scale bar is 20 μm. (**c**) The three panels show the distribution of power density on the taper surface when 12 mW/mm^2^ are emitted for three input angles, namely θ_in_=15°,24°,35°. As light is out-coupled from a wider region when the input angle increases, the input power has to be varied accordingly to maintain a constant output power density.

In order to verify that the emission features described so-far are not modified by tissue absorption and scattering, light delivery performances were tested in brain tissue by inserting 0.66 NA TFs in fixed mouse brain slices previously stained with CYBR green (Figure 4a, see Methods). Spatially selective light delivery was characterized by stimulating fluorescence emission from the taper while remotely varying the input angle. Figure 4b shows a false color overlay of the fluorescent emission excited over four adjacent brain regions. As for the characterization in fluorescein, we used the emission profiles measured along the taper surface to extract centroid *c* and FWHM *Δc* of the emission region for each input angle. As shown in figure 4c, scattering from brain tissue slightly widens the distribution of the light emitted for a given modal subset, but the overall behavior of *c vs θ*_*in*_ is not affected, with some deviations from the linearity measured in fluorescein due to the non-homogeneity of tissue scattering.

**Figure 4:**
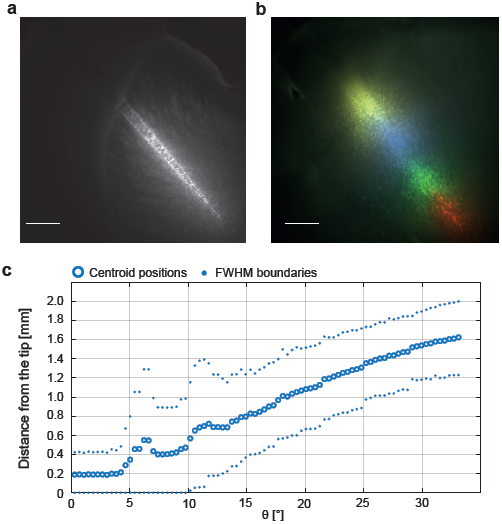
High NA TFs Light delivery in brain tissue. The figure shows a NA=0.66 TF inserted in the striatum region of coronal mouse brain slice. **(a)** Large volume illumination can be obtained by injecting light over the fiber full NA, scale bar is 500 μm. (**b**) False color overlay of adjacent regions illuminated with spatially selective light delivery, scale bar is 500 μm. (**c**) Position of the intensity centroid of the emission region (blue circles) and positions of the FWHM values of the emission region (pale blue dots) with respect to the input angle.

### 2.3 Yellow and multi-color light delivery with high-NA TFs

New optogenetic actuators have been developed in the last years, with activation peaks spanning the visible spectrum**^24–26^**. In particular, optogenetic inhibition of neural activity has been demonstrated by delivering yellow light over neural population transfected with inhibiting opsin probes, such as Halorhodopsin**^26^**. Moreover, a recent work used a TF-based device to achieve neuronal inactivation over a large volume in the non-human primate cortex by targeting a red light-sensitive halorhodopsin (Jaws) with 635 nm light **^23^**. Since both the number of guided modes and the width of the k_t_ values sustained by TFs decrease at higher wavelengths (M∼4×10^5^, 3×10^5^, 2×10^5^ for 473, 561 nm and 635 nm respectively, see equations 2 and 3), it is important to assess TFs performances for site-selective optogenetic inhibition. Fluorescence emission profiles for angle-selective injection (Figure 5a-b) were acquired in a water:eosin solution (see Methods) to test fiber response at 561 nm light. As observed for blue wavelengths, we found that the position of the emission region centroid moves linearly with the input angle along a total region of ∼1.8 mm (figure 5c) (fit slope 60.11 μm/° and RMSE 25 μm). Therefore, TFs spatially-selective modal de-multiplexing properties can be exploited at multiple wavelengths. Figure 5d,e show the independent routing of two laser beams at 473nm and 561nm into a TFs injected in a coronal mouse brain slice co-stained with SYBR-green and MitoTracker red. The false-color fluorescence image in figure 5e was imaged to detect the green and red fluorescence generated by the blue and the yellow light, respectively. It shows that a single device can perform multi-color light deliver over two functionally distinct brain regions: light injected at low input angle is out-coupled in the striatum region (and excites red fluorescence), while blue light, injected at high input angle, is emitted in the deepest layers of cortex (exciting green fluorescence).

**Figure 5:**
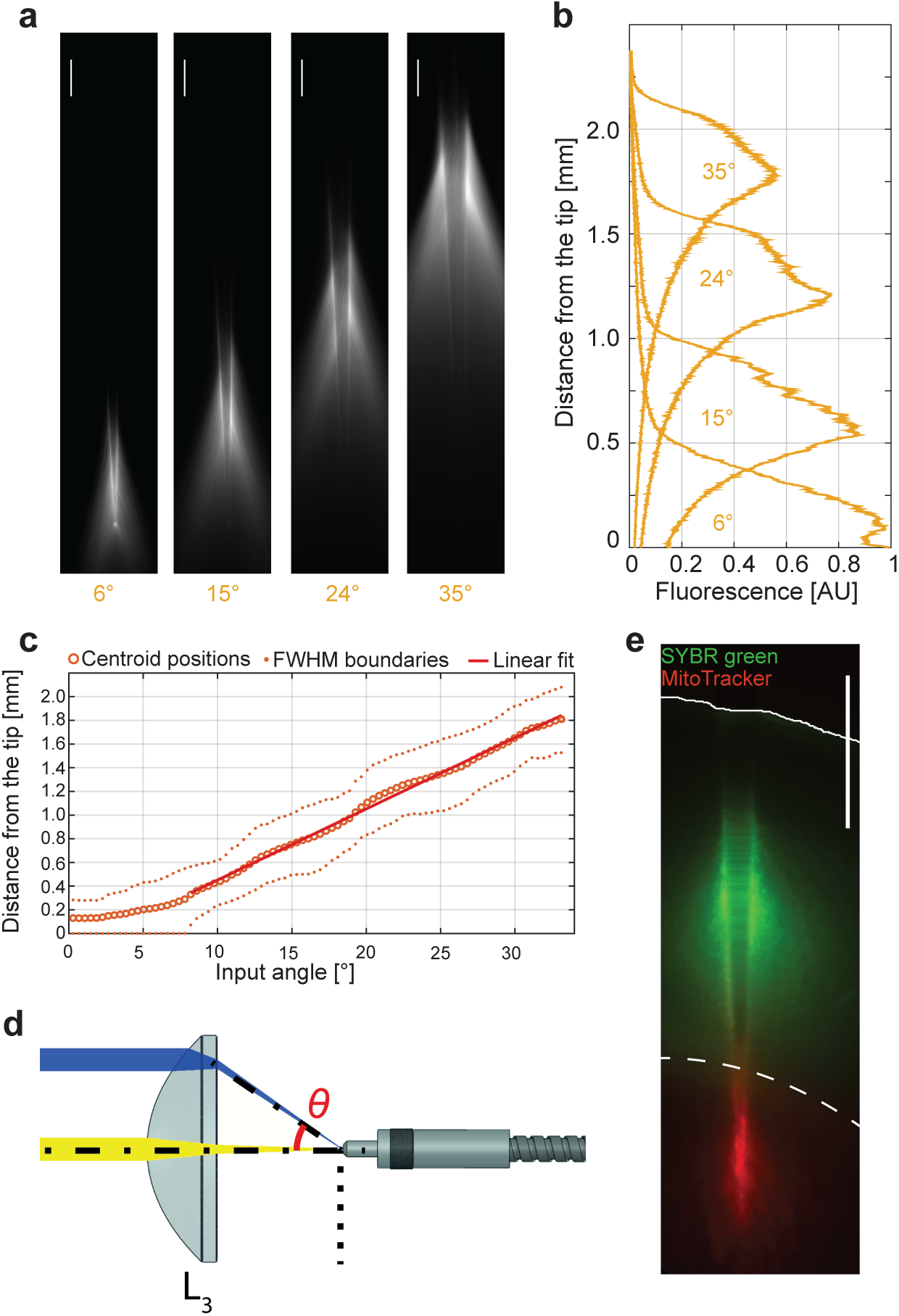
Yellow and two-color light delivery with high-NA TF. (**a**) Spatially selective light delivery observed for a NA=0.66, 200 μm core size fiber injected with yellow (561 nm) light and submerged in a water-eosin solution. Scale bar is 200 μm (**b**) Emission profiles measure along the taper surface for the input angles shown in panel (**a**). **(c**) Position of the intensity centroid of the emission region (orange circles) and positions of the FWHM values of the emission region (pale orange dots) with respect to the input angle. The red line indicates a linear fit on the centroid position. (**d**) Independent injection of yellow and blue light exciting different modal subset with a single lens (**e**) False color overlay showing a simultaneous two-color light delivery with a NA=0.66 TF inserted in a mouse brain coronal slice co-stained with SYBR green and Mito Tracker. Scale bar is 750 μm.

## 3. Discussion

Delivering light in a controlled fashion is required in order to fully exploit the potential of optogenetic techniques in controlling and monitoring neural circuits ^29^. To this end, light delivery based on TFs offers an innovative tool to reach deep brain regions that are precluded to ordinary optogenetic stimulation protocols. This study examines the link between mode subsets supported by the waveguide and TFs emission geometries, and describes how constitutive parameters of the waveguide allow acting on the total number of guided modes and on the related transversal component of the propagation vector. This resulted in the possibility of tailoring emission length of TFs devices from ∼0.4 mm to ∼3 mm. By injecting defined subsets of guided modes within specific k_t_ intervals, far field analysis allowed measurement of the direct relationship between the light output position along the taper and the injected k_t_ values. This revealed a linear dependence for input angles θ_in_>10° for both blue and yellow light. Within the linear region, the output power remains almost constant, while the emitted power density slightly decreases due to the increase of waveguide diameter when the emitting region moves farther from the taper tip.

TFs represent a unique approach to deliver light into the living brain with reduced invasiveness, as they allow for site-selective stimulation over wide volumes without the implantation of electric devices or of multiple waveguides. All these features let us envision that the here-presented results can help neuroscientists in improving experimental protocols targeting sub-cortical regions of the mouse brain, functional structures of larger rodents and non-human primates, as well as analysis methods not compatible with conducting metals (such as opto-fMRI), placing TFs as a great complement to available methods to deliver light into brain.

## Materials and methods

### Device structure and fabrication process

Tapered optical fibers with taper angles ranging from 2° to 8°, fabricated from fiber cords with NA=0.22 core/cladding=50/125 μm, NA=0.39 core/cladding=200/225 μm, NA=0.66 core/cladding=200/230 μm were obtained from OptogeniX (www.optogenix.com). Tapers were fabricated following the previously described procedure**^30^** and connected to ceramic or metallic ferrules with a diameter of 1.25mm.

### Emission properties characterization

#### Data acquisition

The emission properties of the devices were characterized with two different light coupling systems: *(i)* injection of the whole numerical aperture accepted by the fiber and *(ii)* injection of a defined input angle *θ*. Laser light was injected at 473 nm (Laser Quantum Ciel 473) and 561 nm (Coherent OBIS 561nm LS). In both systems, the taper was coupled to a patch fiber with matching NA and core size by a ferrule to ferrule butt-coupling. In the full NA injection configuration of case (*i*) light was coupled to the fiber patch cord with an Olympus objective AMEP-4625 (focal length 4.5 mm, N.A. = 0.65), or with fiber-ports (Thorlabs PAF-SMA-5-A focal length 4.6 mm, N.A. = 0.47, PAF-SMA-7-A focal length 7.5 mm, N.A. = 0.29). To fill the entire clear aperture of the coupling lens, the laser beam was expanded by a proper factor through a beam expander realized with off-the-shelf lenses.

The angle-selective launch system of case *(ii)* was implemented using a galvanometric-mirror based scanning system. Two lenses L1 and L2 (Thorlabs LA1050-A with focal length *f*_1_ = 100 mm and AL50100-A with focal length *f*_2_ = 100 mm, respectively) relayed the laser beam and converted the angular deflection of the GM mirror (Sutter RESSCANGEN) into a displacement *t* perpendicular to the optical axis of the system. Lens L3 (Thorlabs AL4532-A with focal length *f*_*3*_ ^(c)^ = 32 mm and a 45 mm aperture) focused the light onto the patch fiber core. The actual input angle *θ* was measured by registering the displacement of the laser spot on a digital camera placed in front of the coupling lens for a known mirror deflection. The tapered fibers were immersed in a PBS:fluorescein or water:eosin solution to image light emission patterns with input wavelengths of 473 nm or 561 nm, respectively. Images were acquired using an upright epi-fluorescence upright microscope (Scientifica Slicescope) equipped with a 4X objective (Olympus XLFLUOR4X/340 with immersion cap XL-CAP) and a sCMOS camera (Hamamatsu ORCAFlash4.0 V2). Optical output power was measured in air with a power meter (Thorlabs PM100USB with S120VC sensor head). Power coupling efficiency was measured as the ratio between taper and patch fiber optical power output.

Tapered fiber emission in brain slices was measured by inserting the light emitting region of the taper in coronal mouse brain slices fixed in PFA and stained with SYBR green. Excitation laser light at 473 nm was coupled with system (*i*) and (*ii*) into the tapered fiber. Detection of fluorescence was performed through an epi-fluorescence upright microscope (Scientifica Slicescope) equipped with a 4X objective (Olympus XLFLUOR4X/340 with immersion cap XL-CAP) and a sCMOS camera (Hamamatsu ORCA-Flash4.0 V2).

#### Data analysis

A set of intensity profiles was extracted from the fluorescence images collected with coupling scheme *(i)* and *(ii)*. Profiles were measured along a line drawn just outside the taper, parallel to the waveguide surface. Profiles given by full N.A. excitation were used to quantify both the First Emission Diameter (FED) and the total length of the emission region (Emission Length, EL). The FED was defined as the position at which the intensity recorded by the sCMOS sensor is half of the average intensity detected in the pixel with a recorded intensity exceeding 90% of the maximum recorded intensity (supplementary figure 2). The same intensity profiles were used to estimate the power density distribution along the taper when the coupling scheme *(ii)* was used. Assuming a rotational symmetry around the taper axis of the power density distribution *p*, a Matlab code has been developed to calculate *p* starting from the total output power and the intensity profile extracted from the image.

Spatial selectivity in light out-coupling from the taper was quantified from the profiles measured with angle-selective excitation. The centroid and Full Width at Half Maximum (FWHM) positions of the emission region were measured for each input angle *θ* with respect to the fiber tip.

Linear fits as function of the input angle *θ* were computed on the centroid position and both the FWHM boundaries positions. Data points for which the lower bound of the emission region was coincident with the fiber tip were excluded from the fit computation because the presence of the taper tip affects these measurements by breaking the shape symmetry.

### Injected *k*_*t*_ measurements

The value of the transversal propagation constant *k*_t_ of the light guided by the optical fiber as a function of the input angle was measured by imaging the farfield pattern of the light outcoupled by the facet of an optical fiber stub while changing *θ* at the input facet, as previously described^22,31^. In particular, Olympus objective AMEP-4625 (focal length 4.5 mm, N.A. = 0.65) was used to generate the farfield pattern, and a two-lens imaging system (realized with Thorlabs AC254-040-AML and AL50100-A) was used to magnify the pattern over the chip of the sCMOS sensor (Hamamatsu ORCA-Flash4.0 v2). The signal detected at distance *r* from the center of the chip is associated to *k*_t_ by the equation

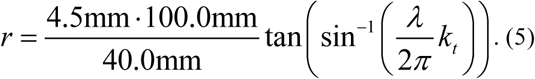

For each input angle, the k_t_ at which the maximum signal is detected and the half-prominence width of the peak were extracted from the disk- or ring-shaped images recorded by the sCMOS sensor.

### Ray tracing simulations

Ray tracing simulations were performed with the commercial optical ray tracing software Zemax-OpticStudio (http://zemax.com). TFs were modeled as straight core/cladding nested cylinders followed by a conical taper section. Core/cladding refractive indexes were assumed homogeneous and set as retrieved from online database**^32^**. In detail, core refractive index was set as 1.6343 and cladding one was set as 1.4900. Refractive index in the tapered region was assumed homogeneous and simulated as the average between core and cladding refractive indexes weighted on the respective cross sectional areas. In particular, the assumed taper refractive index was 1.62. Core/cladding diameters were set as the manufacturer nominal values of 200/230 μm. The length of the core/cladding segment was set to 50 mm whereas the length of the taper is a function of the taper angle.

For full NA light injection simulation, the source was modeled as a circular homogeneous light distribution filling an ideal paraxial lens. The lens NA was modeled in accordance with the injected fiber NA. For selective angle injection simulation, the source was modeled as a circular homogeneous light distribution coupled in the fiber with an ideal paraxial lens. Different input angles were simulated by modulating the tilt of the source-lens system with respect to the fiber axis.

The irradiance profiles of the emitted light were detected by simulating a rectangular, pixelated detector positioned in close proximity of the taper surface. The detector length matched the taper length. Ray tracing simulations were performed launching 5M rays.

## Additional information

Potential competing financial interests: M.D.V., F. Pisanello, B.L.S., and L.S. are cofounders of Optogenix LLC, a company based in Italy that produces and markets the tapered fibers described here.

## Acknowledgments

The authors thank Andrea Della Patria for providing guidance and useful insights with Ray Tracing simulations. F. Pisanello, F. Pisano, E.B. and E.M acknowledge funding from the European Research Council under the European Union’s Horizon 2020 research and innovation program (#677683); B.S. and M.D.V. acknowledge funding from the European Research Council under the European Union’s Horizon 2020 research and innovation program (#692943). L.S., M.P., M.D.V. and B.L.S. are funded by the US National Institutes of Health (U01NS094190); G.M. and B.L.S. are funded by the Simons Collaboration on the Global Brain.

---

**Supplementary figure 1:**
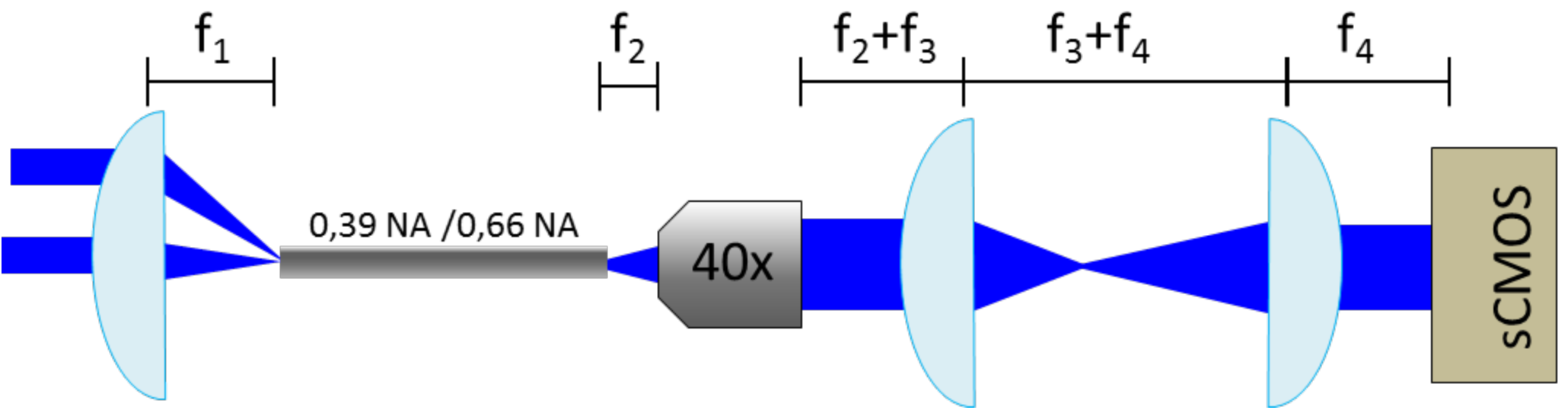
Optical system for far-field imaging of the fiber output. Light is injected into the fiber with an aspheric NA=0.61 lens (f1=32 mm). Light emitted from the fiber is collected with a 40x objective (f2=4.5 mm) and relayed on a sCMOS chip with an achromatic doublet, f3=40 mm, and a 50 mm, f4=100 mm lens.

**Supplementary figure 2:**
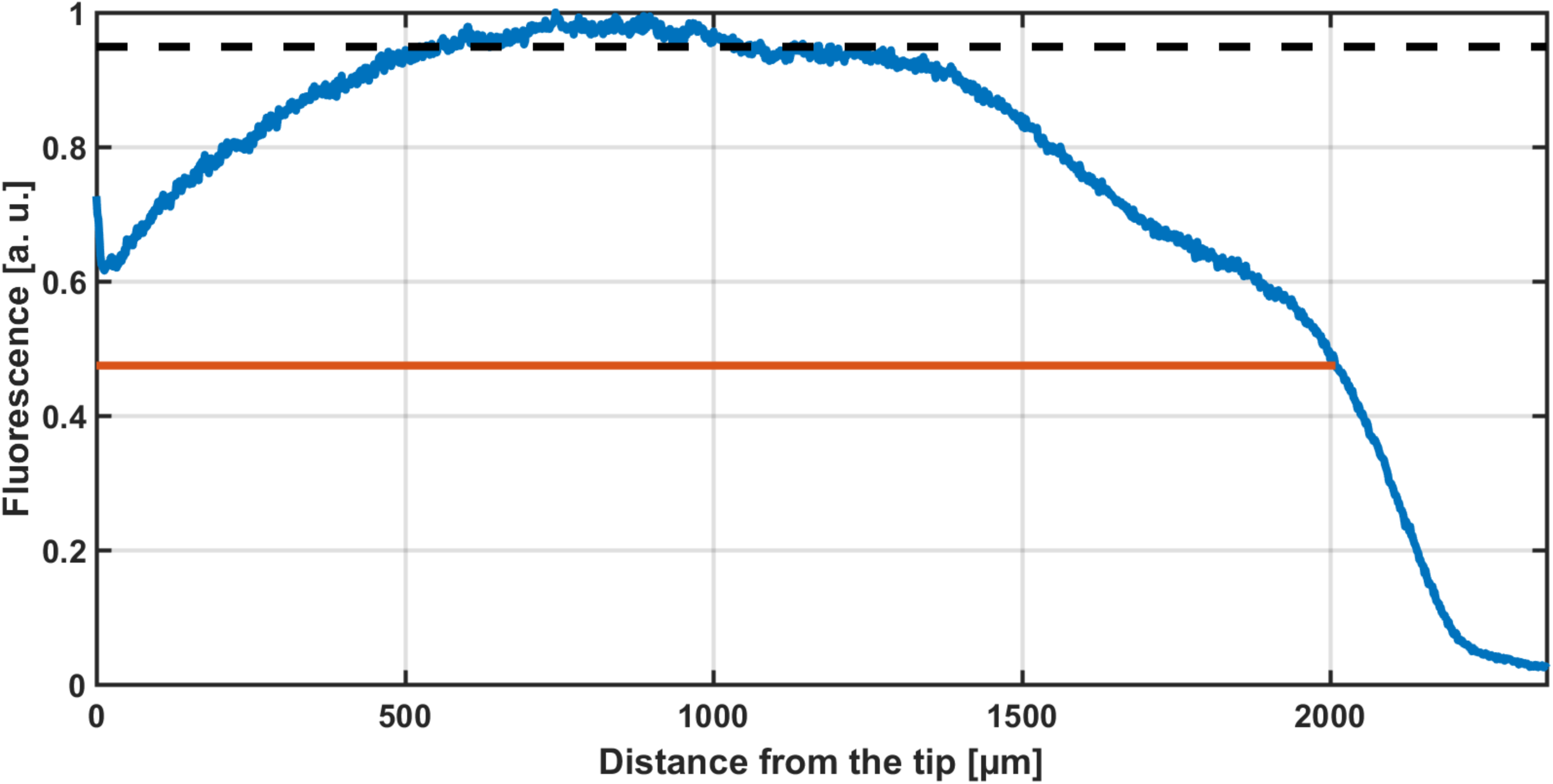
Definition of the First Emission Diameter. The First Emission Diameter is defined as the waveguide diameter at the taper section at which light intensity recorded by the sCMOS sensor (blue curve) fall below half of the value indicated by the black dashed line, representing the average intensity of the datapoints exceeding 90% of the maximum recorded intensity. The red line highlights the region between the taper tip and the FED.

